# Antibiotics that affect translation can antagonize phage infectivity by interfering with the deployment of counter-defences

**DOI:** 10.1101/2022.09.07.506948

**Authors:** Benoit J. Pons, Tatiana Dimitriu, Edze R. Westra, Stineke van Houte

**Affiliations:** ESI, Biosciences, University of Exeter, TR10 9FE Penryn, UK

## Abstract

It is becoming increasingly clear that antibiotics can both positively and negatively impact the infectivity of bacteriophages (phage), but the underlying mechanisms often remain unclear. Here we demonstrate that antibiotics that target the protein translation machinery can fundamentally alter the outcome of bacteria-phage interactions by interfering with the production of phage-encoded counter-defence proteins. Specifically, using *Pseudomonas aeruginosa* PA14 and phage DMS3vir as a model, we show that bacteria with CRISPR-Cas immune systems have elevated levels of immunity against phage that encode anti-CRISPR (*acr*) genes when translation inhibitors are present in the environment. CRISPR-Cas are highly prevalent defence systems that enable bacteria to detect and destroy phage genomes in a sequence-specific manner. In response, many phages encode *acr* genes that are expressed immediately following infection to inhibit key steps of the CRISPR-Cas immune response. Our data show that while phage carrying *acr* genes can amplify efficiently on bacteria with CRISPR-Cas immune systems in the absence of antibiotics, the presence of antibiotics that act on protein translation prevents phage amplification, while protecting bacteria from lysis. These results help to understand how antibiotics-phage synergy and antagonism depend on the molecular interactions that define phage infectivity and host immunity.

## Introduction

In natural environments, phages are estimated to be 10 times more abundant than bacteria, and to cause the lysis of 20 to 40 % of the bacterial biomass each day^1^. To resist against phage infection, bacteria have evolved a wide range of defence mechanisms^2–4^ organised in several ‘lines of defence’, providing immunity against a wide variety of viruses^5,6^. Among these defence mechanisms, the CRISPR-Cas (Clustered Regularly Interspaced Short Palindromic Repeat, CRISPR associated) system provides acquired immunity against previously encountered phages and other mobile genetic elements^7–10^. It relies on acquisition of fragments from invading phage genetic material (spacers) from earlier failed infections, which are inserted in a specific CRISPR locus on the bacterial chromosome. CRISPR RNAs (crRNAs) are then transcribed and processed from the CRISPR loci and form a surveillance complex with Cas protein(s). Guided by the crRNA, this surveillance complex recognizes sequences matching the spacer (protospacers) in invading genetic material, leading to sequence-specific cleavage of the invader. CRISPR-Cas systems are found in approximately 40% of bacterial genomes, making them one of the most prevalent defence systems identified so far, and are classified in 2 classes, 6 types and 33 subtypes based on differences in number and nature of associated proteins^10^.

Phages are not defenceless against CRISPR-Cas systems as they have evolved counter-defence mechanisms during their struggle against their bacterial foes^3,11^. Some phages encode small peptides, called anti-CRISPR proteins (Acr), that hinder binding or cleavage of the phage genome by the CRISPR-Cas system^12–14^. Acrs are not *a priori* present in the phage particles but are expressed only at the start of the infection, at very high levels^15,16^. When faced with a CRISPR-immune host (i.e., that has a fully matching spacer targeting the phage), Acr production is not always fast enough to completely inactivate the surveillance machinery, leading to cleavage of the phage genetic material^17,18^. However, despite phage cleavage, the Acr protein produced will leave bacteria in an immunosuppressed state, in which part of the surveillance complexes are inhibited by Acr, thus increasing the probability that a subsequently infecting phage can successfully replicate on this immunosuppressed host ^17–19^.

Recent work has shown that the presence of bacteriostatic antibiotics (antibiotics inhibiting cell growth without killing) favour acquisition of CRISPR-Cas immunity during infection with phages that lack Acr activity^20^. Sub-inhibitory doses of bacteriostatic antibiotics slow down both bacterial growth and phage replication, thereby lengthening the phage replication cycle and hence allowing more time for bacteria to acquire new spacers against the phage before being lysed. Phage-antibiotic interactions can range from synergy^21,22^ to antagonism^23^. While often a mechanistic explanation for the observed interaction is lacking (^24^ and references herein), several different mechanisms have been identified, including antagonism caused by a decreased host RNA synthesis^25,26^ and synergy mediated by phage-mediated impairment of antibiotic resistance development coupled by an antibiotic-mediated hindrance of phage resistance apparition^27^. The paper by Dimitriu *et al* identifies a previously unknown mechanism driving antagonism between bacteriostatic antibiotics and phages that infect bacteria carrying a functional CRISPR-Cas system.

Antibiotics impair bacterial growth or kill cells by acting on various molecular targets^28^. One of the most common targets is the ribosome, an enzymatic complex responsible for the translation of messenger RNA into functional polypeptides, and hence essential for bacterial survival and growth. Antibiotic compounds from a wide variety of classes bind to only a few sites in the 30S or 50S ribosomal subunits, which disturbs initiation, elongation, or termination of translation, even at sub-inhibitory antibiotic doses^29–32^.

Since successful Acr-phage amplification relies on the strong production of Acrs at the onset of infection, we hypothesised that translation inhibitor antibiotics have the potential to interfere with Acr-induced immunosuppression, thereby effectively re-sensitising the phage to full CRISPR-Cas immunity. Using the *Pseudomonas aeruginosa* strain PA14 and its lytic phage DMS3*vir* as a model system, we show that sub-inhibitory doses of translation inhibitory antibiotics block Acr-induced immunosuppression and hence successful phage replication. Concomitantly, infected bacteria benefit from the presence of these antibiotics, suggesting an antagonistic interaction between translation inhibitor antibiotics and Acr-phages infecting CRISPR-immune bacteria.

## Results

### Translation inhibitors antibiotics disrupt Acr-mediated inhibition of CRISPR-Cas immunity

The strong and early production of Acrs^15,16^ during phage infection suggests that their efficiency might be impaired in the presence of translation inhibitor antibiotics. To test the hypothesis that these antibiotics may impact the effectiveness of Acr proteins during phage infection, we infected *P. aeruginosa* UCBPP-PA14 (PA14) with the lytic phage *DMS3mvir-* AcrIF1, which carries a type I-F *acr* gene^33^, or the isogenic control phage DMS3*mvir*, which does not carry a type I-F *acr*. Prior to infection, bacteria were grown for 2 hours in the presence of minimum inhibitory concentration (MIC) or sub-MIC doses of five antibiotics belonging to different chemical classes (Table S1). Four of these antibiotics (chloramphenicol, Chl, erythromycin, Ery, tetracycline, Tet and gentamycin, Gm) target protein translation, while one (carbenicillin, Carb) has no effect on translation^20,29^. Wild type bacteria, which carry a type I-F CRISPR-Cas immune system that targets phage *DMSmvir* and DMS3*mvir*-AcrIF1 *a priori* (CRISPR-immune), or an isogenic CRISPR-knockout (CRISPR-KO) control strain were infected at low multiplicity of infection (MOI) with either DMS3*mvir* or DMS3*mvir*-AcrIF1. After washing away antibiotics and phage, we assessed the relative transformation efficiency (RTE) of the bacteria by transforming them with a plasmid either targeted (T) or not targeted (NT) by the PA14 CRISPR-Cas system (Figure 1).

**Figure 1.**
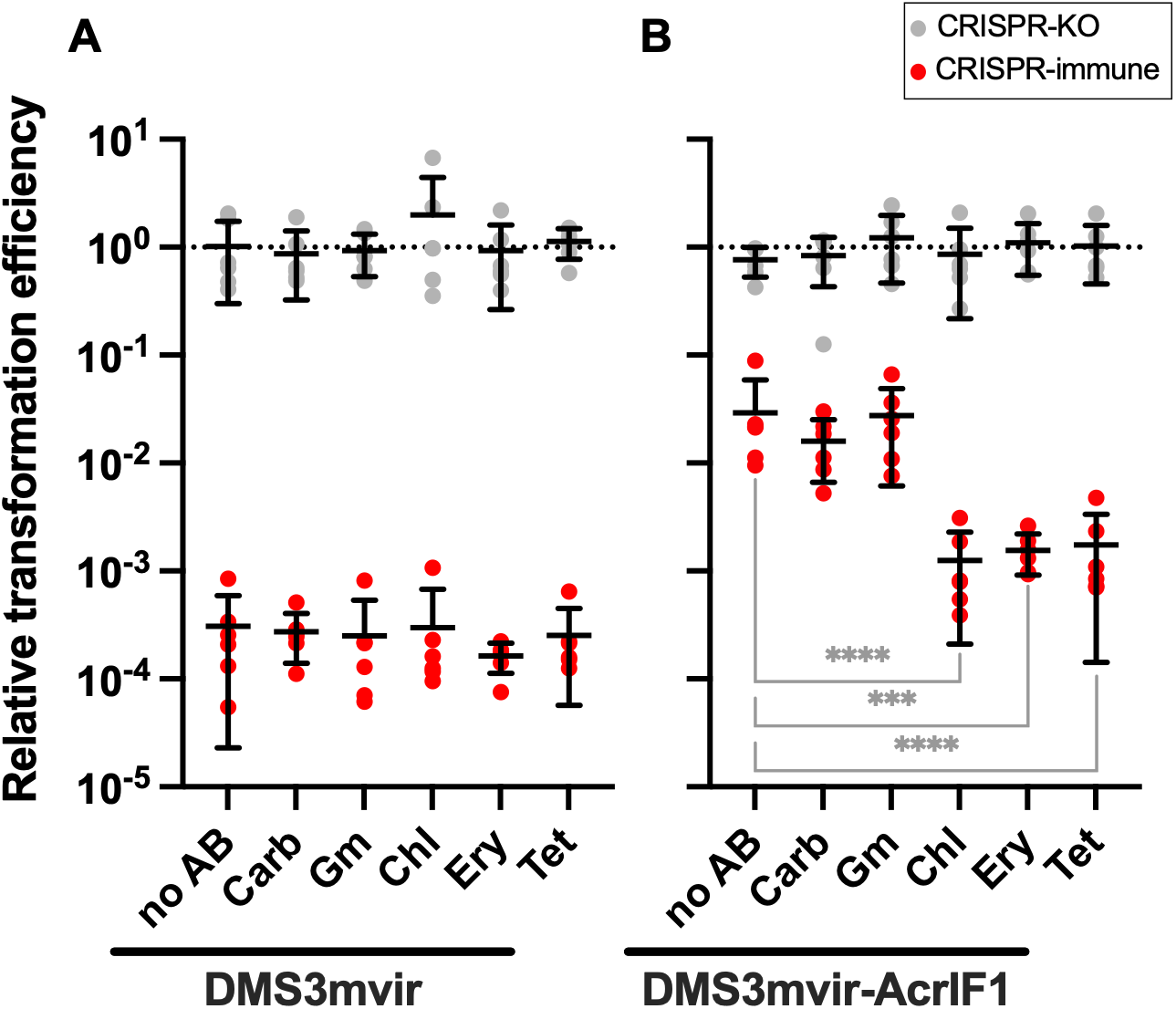
Bacteriostatic translation inhibitors disrupt Acr-mediated inhibition of the CRISPR-system. Relative transformation efficiencies (targeted plasmid/non-targeted plasmid) of PA14 CRISPR-KO (grey data points) or CRISPR-immune (red data points) pre-infected with phage *DMS3mvir* (A) or DMS3*mvir*-AcrIF1 (B), in the absence (no AB) or presence of different antibiotics (see Table S1). Each data point represents an independent biological replicate (n = 6), and the mean ± standard deviation for each treatment is displayed as black bars. Asterisks show treatments that are different from the no-antibiotic control (Dunnett, *** 0.0001<p<0.001, **** p<0.0001).

Neither the phage treatment nor the antibiotic treatment had influence on the RTE of the CRISPR-KO strain (Figure 1). As previously shown^18^, the CRISPR-immune strain infected with DMS3*mvir*-AcrIF1 displayed a higher RTE than when infected with DMS3*mvir* due to lasting immunosuppression by the Acr. None of the antibiotic treatments affected the RTE of the CRISPR-immune strain infected with DMS3*mvir* compared to the no-antibiotic control (p>0.9 for all antibiotics) (Figure 1A). In contrast, treatment with Chl, Ery and Tet significantly decreased the RTE when the CRISPR-immune strain was infected with DMS3*mvir*-AcrIF1 compared to the no-antibiotic control (p<0.0001, p=0.0004, p<0.0001, respectively; Figure 1B). Conversely, Carb and Gm did not affect immunosuppression in CRISPR-immune cells by DMS3*mvir*-AcrIF1 (p=0.80 and p>0.99). Overall, these results suggest that Chl, Ery and Tet, but not Gm or Carb can block immunosuppression induced by AcrIF1.

### Translation inhibitor antibiotics decrease infection efficiency of Acr-phage

Based on the observation that some translation inhibitors interfere with Acr-induced immunosuppression, we then hypothesised that these antibiotics would affect DMS3*mvir*-AcrIF1 replication in a CRISPR-immune host. To test this, we infected CRISPR-KO and CRISPR-immune cells with DMS3*mvir* or DMS3*mvir*-AcrIF1 at MOI=1 in the presence or absence of antibiotics and measured phage titres after 24h (Figure 2). Control experiments with a CRISPR-KO strain showed that antibiotics have an impact on phage amplification, with the largest effect size for Gm (Figure 2A). Crucially, there was no difference between phage DMS3*mvir* and DMS3*mvir*-AcrIF1. In contrast, in CRISPR-immune bacteria, phages were only able to amplify above input titre when carrying the *acrIF1* gene (Figure 2B). Moreover, in the presence of any of the four translation inhibitors, DMS3*mvir*-AcrIF1 was unable to amplify (p<0.0001 for all treatments, compared with the no-antibiotic control), whereas Carb did not interfere with phage amplification (p>0.99). We also noted that in the case of Gm, a decline in phage titre was observed in the absence of CRISPR-Cas, and this was independent of the presence or absence of the phage *acrIF1* gene (Figure 2A). This suggests that Gm causes an overall phage fitness decrease, in line with previous studies showing that aminoglycosides can inhibit phage infectivity^23,34,35^. Thus, these results show that the three translation inhibitors Chl, Ery and Tet interfere with the ability of DMS3*mvir*-AcrIF1 to block CRISPR-Cas immunity.

**Figure 2.**
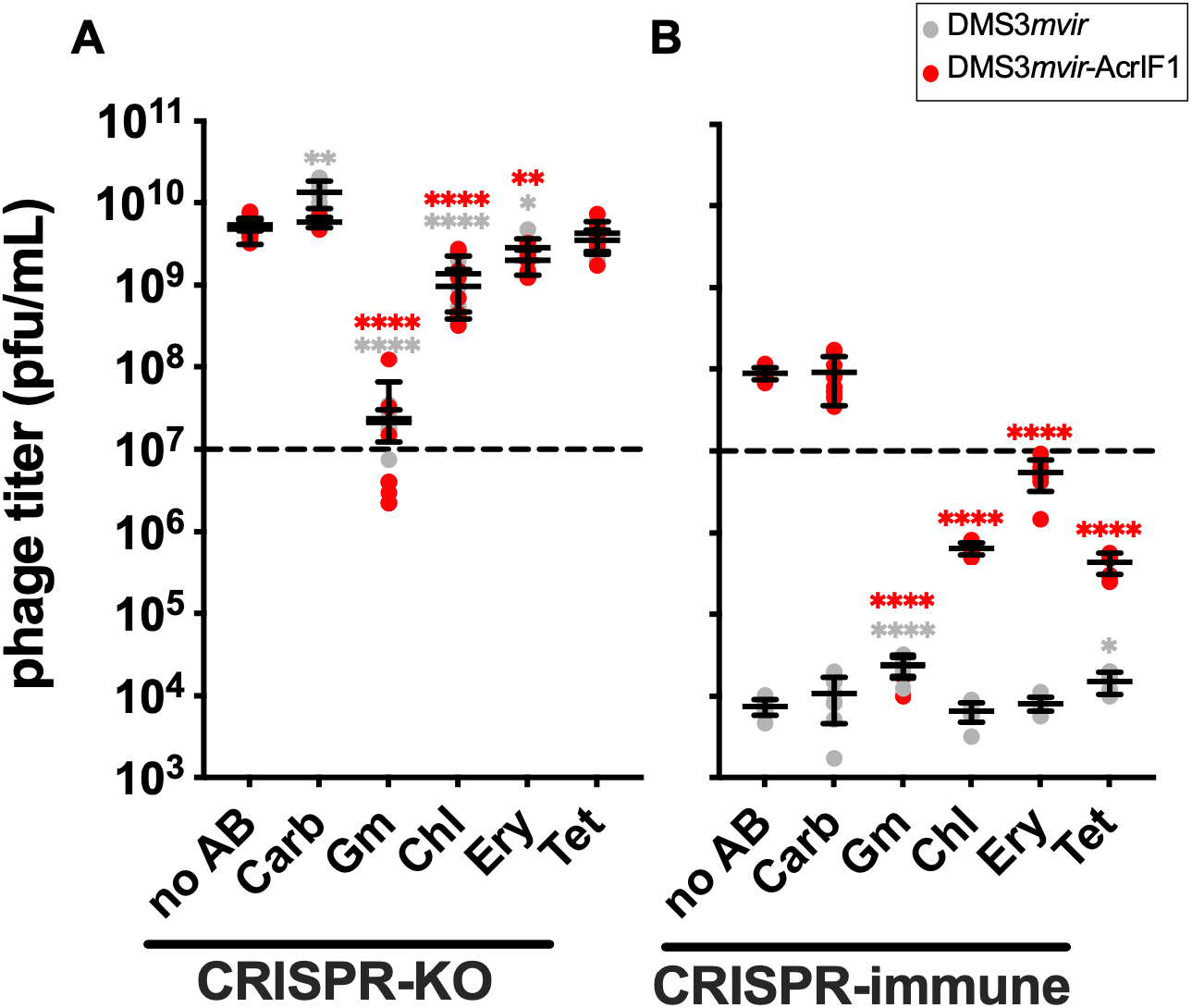
Translation inhibitor antibiotics decrease infection efficiency of Acr-phage. Effects of different antibiotic treatments (see Table S1) on DMS3*mvir* (grey data points) or DMS3*mvir*-AcrIF1 (red data points) titre after 24h of infection on PA14 CRISPR-KO (A) or CRISPR-immune cells (B). The dashed line indicates the phage titre at t=0h. Each data point represents an independent biological replicate (n = 8) with limit of detection at 250 plaque forming units (PFUs)/mL, and the mean ± standard deviation for each treatment is displayed as black bars. Asterisks show treatments that are different from the no-antibiotic control (Dunnett, * 0.01<p<0.05, ** 0.001<p<0.01, *** 0.0001<p<0.001, **** p<0.0001).

### Translation inhibitor antibiotics protect CRISPR-immune cells from Acr-phages

Since Chl, Ery and Tet hinder DMS3*mvir*-AcrIF1 amplification on CRISPR-immune bacteria, we predicted that these antibiotics would protect infected cells from phage-induced lysis. We thus evaluated the impact of antibiotics on the optical density at λ=600nm (OD_600_) of PA14 CRISPR-KO or CRISPR-immune after 24h of infection by either DMS3*mvir* or DMS3*mvir*-AcrIF1 at MOI=1 (Figure 3). As expected, for each antibiotic treatment, bacterial growth of CRISPR-KO cells following phage infection was independent of the presence or absence of phage carrying *acrIF1* (Figure 3A). Conversely, in the absence of antibiotics, the OD_600_ of CRISPR-immune cells was higher following infection with DMS3*mvir* than DMS3*mvir*-AcrIF1, and the presence of Carb did not affect this pattern (Figure 3B). However, the presence of Chl, Ery and Tet allowed CRISPR-immune cells to grow to similar OD_600_ when infected by DMS3*mvir* and DMS3*mvir*-AcrIF1, thus removing any effect of phage-encoded *acrIF1* on host growth. Treatment with Gm resulted in levels of bacterial growth similar to the no-phage controls (Figure S1), independent of the presence of a functional CRISPR system in the host, and both in the presence DMS3*mvir* and DMS3*mvir*-AcrIF1 (Figure 3), further supporting a direct effect of this antibiotic on phage infectivity. Overall, these results suggest that Chl, Ery and Tet increase the ability of CRISPR-immune bacteria to resist lysis by Acr-phage, causing phage-antibiotics antagonism in those instances.

**Figure 3.**
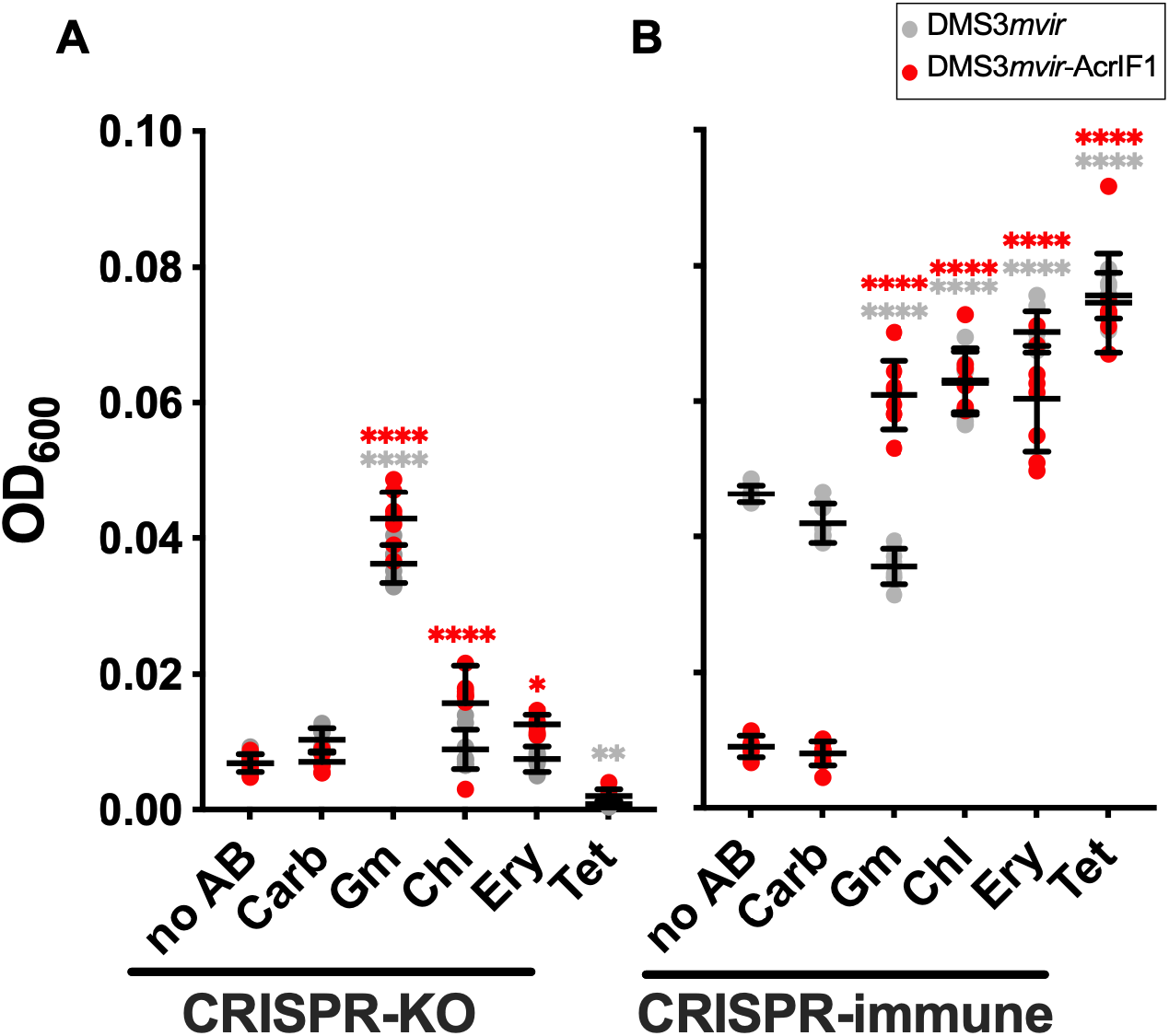
Translation inhibitor antibiotics protect CRISPR-immune cells from Acr-mediated lysis. Effects of the different antibiotic treatments (see Table S1) on bacterial OD_600_ of PA14 CRISPR-KO (A) or CRISPR-immune (B) after 24h of infection by DMS3*mvir* (grey data points) or DMS3*mvir*-AcrIF1 (red data points). Each data point represents an independent biological replicate (n = 8), and the mean ± standard deviation for each treatment is displayed as black bars. Asterisks show treatments that are different from the no-antibiotic control (Dunnett, * 0.01<p<0.05, ** 0.001<p<0.01**** p<0.0001).

## Discussion

In Mu-like phages, such as DMS3, *acr* genes are expressed before genes classified as early expressed, including the transposase^15,36^. Evidence that some Acr-carrying phages need several infections of the same cell to successfully overcome CRISPR-mediated immunity^17,18^ suggest that Acr protein levels, and thus its production, are critical to its ability to overcome CRISPR-Cas immunity. We consequently hypothesised that disturbance in Acr production might affect the outcome of Acr-phage infection on a CRISPR-immune host. More specifically, we propose that antibiotics inhibiting protein translation disturb Acr production and thus interfere with the effectiveness of Acr-phages infecting CRISPR-immune bacteria.

We show here that Acr activity is indeed reduced when CRISPR-immune cells were coexposed to Acr-phages and the translation inhibiting antibiotics Chl, Ery or Tet (Figure 1). Exposure to these antibiotics significantly reduced Acr-phage titre (Figure 2) and allowed CRISPR-immune bacteria to grow to substantially higher density than without antibiotics (Figure 3). Consistent with our hypothesis, these translation inhibitor antibiotics hindered Acr-mediated immunosuppression, presumably through their ability to disturb protein production, even at sub-MIC doses^37–39^. This led to disturbed phage replication, thus providing protection to CRISPR-immune bacteria against Acr-phages.

Despite having no effect on Acr immunosuppression activity, Gm had a similar impact on phage titre and bacterial growth than the previous three antibiotics. However, these effects on phage and bacteria concentration were observed for both phage with and without *acrIF1* and for both CRISPR-immune and CRISPR-KO cells. This suggests that Gm interferes with phage amplification in a way that is independent of CRISPR-Cas and Acr, which results in a lesser impact on bacterial growth. Accordingly, the aminoglycosides antibiotic family, to which Gm belongs, has previously been reported to hinder phage production and favour bacterial growth when used at doses close or above the MIC ^23,34,35^.

Owing to the sharp rise in infections by antibiotic resistant bacterial strains and the health burden that they impose^40^, phage treatment is once again seen as a future replacement to chemical antibacterial treatments^41^. However, bacteria often evolve resistance to phages quickly and effectively through a range of resistance mechanisms, thereby obliterating any therapeutic effect^2,3,42^. Combining phage and antibiotics therapy has been proposed as a way to circumvent this, by imposing two different selective pressures on bacteria^43^. Such an approach has been studied in *in vitro* models, and is now being applied in clinical trials, showing promising effects in comparison with phage or antibiotic treatments alone (^44,45^ and references herein). Another way to tackle bacterial resistance to phage is the use of natural or engineered phage carrying counter-defence mechanisms. Although this strategy has not yet been tested in clinical use, Acr-phage are now envisioned as a potential phage therapy tool to target CRISPR-immune bacteria^46–49^, and have shown promising results in animal models^50^. However, we show here evidence of a negative interaction between some antibiotics and Acr-phages, if the targeted bacterium is CRISPR-immune to the phage used for therapy. These results, along with previous work showing antagonism between phage and antibiotic treatment (^24^ and references herein), highlight the need to test each phage-antibiotic combination as well as the individual treatments, to evaluate potential synergy or antagonism between them. In addition, this negative interaction between translation inhibitor antibiotics and phages might also disrupt phage-host interactions in non-targeted bacterial species. Given the suggested role of phage in structuring and stabilising microbial communities, such as the gut microbiota^51,52^, disturbing the phage-bacteria interaction network through the use of translation inhibitor antibiotics could therefore have important downstream consequences for human or animal health.

Both biotic and abiotic complexity are known to impact phage-host interactions and their coevolution^53^, and the experimental setting used here is necessarily simpler than a natural environment or clinical setting. In this study we used antibiotic doses near or below the MIC for *P. aeruginosa*. While antibiotics are usually used in high-enough doses to cause lethality, the effective concentration can considerably vary between body compartments, potentially reaching sublethal concentrations^54–57^.

Previous results, focusing on antibiotic effects on the outcome of the battle between phage and hosts carrying CRISPR-Cas immunity, showed that bacteriostatic antibiotics tip the balance in favour of bacteria by slowing down bacterial and phage replication, and hence allowing more time for bacteria to acquire spacers against invading phages^20^. We report here another negative effect on phage by Chl, Ery and Tet, which are also bacteriostatic antibiotics. This suggests that their impact on phage infectivity could be two-fold, by first favouring acquisition of spacers against phages (as bacteriostatic antibiotics) and then by decreasing the efficiency of phage counter-defence against bacterial CRISPR-Cas system (as translation inhibitors).

## Experimental model details

### Bacterial and viral strains

The strain derived from UCBPP-PA14 (PA14) of *Pseudomonas aeruginosa* carrying 2 spacers targeting the phage DMS3*vir* (CRISPR-immune) and the strain UCBPP-PA14 *csy3::LacZ* (CRISPR-KO) with a non-functional CRISPR system were described in ^18^. Bacteria were cultured at 37°C with 180 rpm shaking in LB or M9 minimal medium (22 mM Na2HPO4; 22 mM KH2PO4; 8.6 mM NaCl; 20 mM NH4Cl; 1 mM MgSO4; 0.1 mM CaCl2) supplemented with 0.2% glucose (M9+glucose).

### Phages

Recombinant lytic phages DMS3*mvir* (derived from phage DMS3*vir* to be targeted by 1 spacer in PA14 and 3 spacers in the CRISPR-immune strain^8^) and DMS3*mvir*-AcrIF1 (expressing Acr protein that blocks the PA14 CRISPR I-F system^33^) were used throughout this study. Phage stocks were obtained from lysates prepared on PA14 CRISPR-KO and stored at 4°C.

## Methods

### Infection assays in liquid medium

All infections assays were conducted in M9+glucose (22 mM Na2HPO4; 22 mM KH2PO4; 8.6 mM NaCl; 20 mM NH4Cl; 1 mM MgSO_4_; 0.1 mM CaCl_2_; 0.2% w/v glucose). Overnight cultures grown in M9+glucose were diluted to 2*10^7^ colony forming units (CFUs)/mL in fresh media. 180 μL of the diluted cells were added to each well of a 96-wells plate and were subsequently treated with 10 μL of fresh media containing the appropriate antibiotic concentration (final antibiotic concentration listed in Table 1) and 10 μL of fresh media containing 2*10^7^ PFUs/mL DMS3*mvir* or DMS3*mvir*-AcrIF1 (MOI=1), or no phage as control. Each treatment was performed in 8 independent biological replicates. After 24h of incubation at 37°C with 180 rpm shaking, final bacterial concentration was determined by measuring the optical density at λ=600nm (OD_600_) in a Varioskan Flash Multimode plate reader. Final phage concentration was determined by titration on a soft agar lawn. Phages were extracted by mixing 100 μL of each infection with 10 μL of chloroform. After thorough mixing by pipetting, cells were harvested by spinning at 3500 rpm for 20 minutes and the supernatant containing phages was recovered. A mixture of 8 mL of molten LB soft agar (0.5%) and 400 μL of CRISPR-KO cells grown overnight in LB was poured on top of a hard LB agar (1.5%) lawn. Serial dilutions of extracted phages were spotted on this dried soft agar plate and plaques were counted after incubation overnight at 37°C.

### CRISPR immunosuppression experiment

The CRISPR immunosuppression protocol (Figure 1) was adapted from Landsberger *et al*., 2018^18^. Overnight cultures of PA14 CRISPR-immune or CRISPR-KO grown in LB (approximately 3*10^10^ CFUs) were either unexposed or exposed to antibiotic sub-MIC concentrations (Table S1). Subsequently, bacteria were either uninfected or infected with 10^10^ PFUs of DMS3*mvir* or DMS3*mvir*-AcrIF1. Each treatment was performed in 6 independent biological replicates. After 2 hours of incubation at 37°C with 180 rpm shaking, cells were harvested by spinning at 3500 rpm for 20 minutes.

The phage titre was quantified by spotting 4 μL of serially diluted supernatant on LB soft agar plates. After incubation overnight at 37°C, plaques were counted.

Bacteria pellets were washed twice in 1 mL of 300 mM sucrose and resuspended in 300 μL of 300 mM sucrose. The resuspended bacteria were divided in three 100 μL samples. One sample was serially diluted and plated on LB agar to enumerate total bacterial CFUs before transformation (in order to verify that all treatments have equal bacterial concentration before transformation) and the other two were electroporated with either plasmid pHERD30T (not targeted, NT) or a pHERD30T-derived plasmid targeted by the PA14 CRISPR-Cas system (targeted, T)^18^. Transformed bacteria were allowed to recover for 1h at 37°C with 180 rpm shaking in 1 mL of LB. After recovery, bacteria were pelleted and resuspended in 100 μL LB, plated on LB agar plates supplemented with 50 μg/mL Gentamycin to select for transformants, and incubated overnight at 37°C.

Relative transformation efficiency (RTE) was calculated for each treatment as (number of colonies transformed with targeted plasmid)/(number of colonies transformed with non-targeted plasmid).

### Quantification and statistical analysis

All statistical analyses (two-way Anova with Dunnett post-hoc test) were done with Graphpad Prism version 9.3.1 and statistical parameters are reported in the figure legends or within the results section.

## Acknowledgement

B.J.P. was supported by funding from the UK Biotechnology and Biological Sciences Research Council (BB/S017674/1) awarded to S.v.H. S.v.H. also acknowledges funding from BBSRC grant BB/R010781/1 and EPSRC grant EP/X026507/1, and E.R.W received grants from the European Research Council under the European Union’s Horizon 2020 research and innovation programme (ERC-2017-ADG-788405 and ERC-STG-2016-714478).

## Author contributions

Conceptualisation, B.J.P., T.D., E.R.W., and S.v.H..; Methodology, investigation, and formal analysis, B.J.P.; Writing – Original Draft, B.J.P.; Writing – Review & Editing, B.J.P., T.D., E.R.W., and S.v.H.; Funding Acquisition and supervision, E.R.W. and S.v.H.

