## Supplemental Figure 1 for "Antibiotics that affect translation can antagonize phage infectivity by interfering with the deployment of counter-defences"

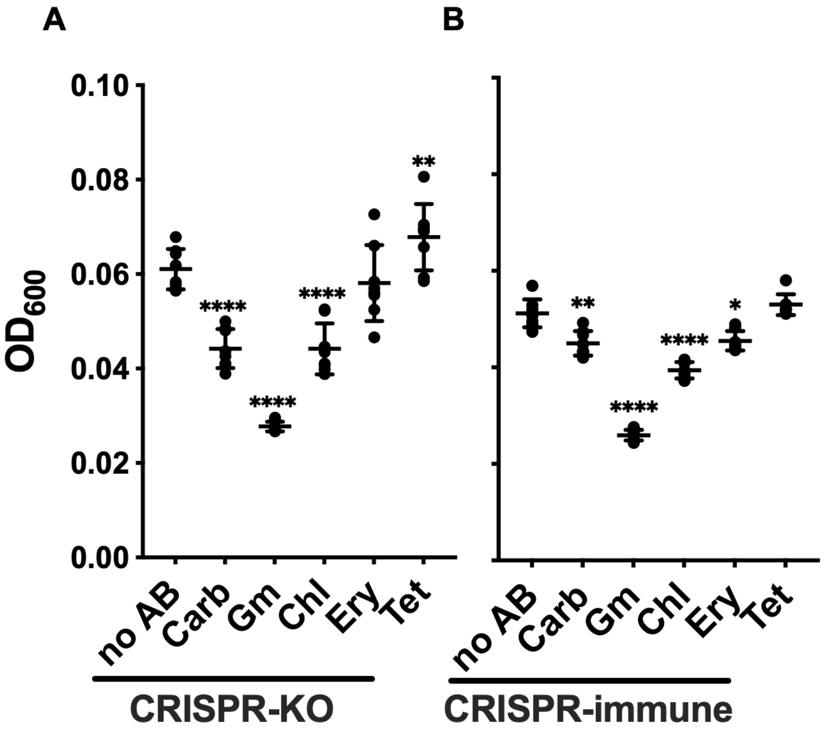


**Figure S1. Translation inhibitor antibiotics do not impact cell growth in the absence of phages.**

Effects of the different antibiotic treatments (see Table S1) on bacterial $OD_{600}$ of PA14 CRISPR-KO (A) or CRISPR-immune (B) after 24h of growth. Each data point represents an independent biological replicate (n = 8), and the mean ± standard deviation for each treatment is displayed as black bars. Asterisks show treatments that are different from the no-antibiotic control (Dunnett, * $0.01<p<0.05$, ** $0.001<p<0.01$**** $p<0.0001$).
