## Supplemental Table 1 for "Antibiotics that affect translation can antagonize phage infectivity by interfering with the deployment of counter-defences"

**Supplementary material**

| Name | | Class | Target | Bacteriostatic/ bactericidal | MIC (µg/mL) | Standard (µg/mL) |
|---|---|---|---|---|---|---|
| Carbenicillin | Carb | Penicillin | Penicillin binding proteins | bactericidal | 25 | 2.5 |
| Gentamycin | Gm | Aminoglycoside | 30S ribosome subunits | bactericidal | 1.25 | 0.63 |
| Chloramphenicol | Chl | Chloramphenicols | 50S ribosome subunits | bacteriostatic | 30 | 25 |
| Erythromycin | Ery | Macrolides | 50S ribosome subunits | bacteriostatic | 100 | 100 |
| Tetracycline | Tet | Tetracyclines | 30S ribosome subunits | bacteriostatic | 10 | 2.5 |

**Table S1. Antibiotics used in this study.**

Antibiotic abbreviations, classes[29], molecular targets[29], overall bacteriostatic/bactericidal[20], minimum inhibitory concentration (MIC, µg/mL)[20] and standard concentrations used (µg/mL) are shown for all antibiotics used in this study.

6
